# Rice Gene Index (RGI): a comprehensive pan-genome database for comparative and functional genomics of Asian rice

**DOI:** 10.1101/2023.02.14.528456

**Authors:** Zhichao Yu, Yongming Chen, Yong Zhou, Yulu Zhang, Mengyuan Li, Yidan Ouyang, Dmytro Chebotarov, Ramil Mauleon, Hu Zhao, Weibo Xie, Kenneth L. McNally, Rod A. Wing, Weilong Guo, Jianwei Zhang

**Affiliations:** National Key Laboratory of Crop Genetic Improvement, Hubei Hongshan Laboratory, Huazhong Agricultural University, Wuhan, 430070, China; Frontiers Science Center for Molecular Design Breeding, Key Laboratory of Crop Heterosis and Utilization (MOE), and Beijing Key Laboratory of Crop Genetic Improvement, China Agricultural University, Beijing 100193, China; Center for Desert Agriculture, Biological and Environmental Sciences & Engineering Division (BESE), King Abdullah University of Science and Technology (KAUST), Thuwal, 23955-6900, Saudi Arabia; International Rice Research Institute (IRRI), Los Baños, 4031 Laguna, Philippines; Arizona Genomics Institute, School of Plant Sciences, University of Arizona, Tucson, Arizona 85721, USA

**Keywords:** Rice, Pan-genome, Homologs, Index, Visualization

## Abstract

To integrate the genomic information of the rice pan-genome, we performed comparative analyses and established a user-friendly Rice Gene Index (RGI, https://riceome.hzau.edu.cn) platform with 16 platinum standard reference genomes and supplementary transcriptome data. To logically organize and scientifically the index of 744,233 genes among rice accessions, we detected 112,658 Ortholog Gene Indices, and provide ‘GeneCard’ pages to query genomic, transcriptomic, and homology information for each gene. The RGI allows users to search for relationships and comprehensive information of genes in keyword-based, sequence-based, and relationship-based ways. Furthermore, users can visualize these relationships at local and global scales corresponding to ‘Microcollinearity’ and ‘Macrocollinearity’ modules.

## Background

Asian rice (*Oryza sativa*) is the staple food for half the world and is a model crop that has been extensively studied. It contributes ~20% of calories to the human diet (Stein et al., 2018). With the increase in global population and rapid changes in climate, rice breeders need to develop new and sustainable cultivars with higher yields, healthier grains, and reduced environmental footprints (Wing et al., 2018). Since the first gold standard reference genome of rice variety Nipponbare has been published (International Rice Genome Sequencing Project., 2005), an increasing number of rice accessions have been sequenced, assembled, and annotated with global efforts. Nowadays, a single reference genome is obviously insufficient to perform the genetic difference analysis for rice accessions (Huang et al., 2021). Therefore, the pan-genome has been proposed as a solution, which allows the discovery of more presence-absence variants, as compared with single reference genome-based studies (Zhao et al., 2018). Over the past years, several databases, such as RAP-db (https://rapdb.dna.affrc.go.jp), RGAP (http://rice.uga.edu), and Gramene (https://www.gramene.org), have long-term served rice genomic research by providing information based on one or a series of individual reference genomes. To integrate and utilize the genomic information of multiple accessions, we performed comparative analyses and established a user-friendly Rice Gene Index (RGI, https://riceome.hzau.edu.cn) platform. RGI is the first gene-based pan-genome database for rice.

### Data collection

To set up a solid foundation for this database, we selected 16 platinum standard reference genomes of rice accessions that represent the major Asian rice subpopulations when K=15 (Song et al., 2021; Stein *et al*., 2018; Zhou et al., 2020), (Figure 1A). Starting with a set of unified *de novo* annotations performed by Gramene (unpublished) of 14 genomes and 4 published annotations including Minghui 63 (MH63), Zhenshan 97 (ZS97), and Nipponbare (RGAP and RAP-db) (Kawahara et al., 2013; Sakai et al., 2013)), we incrementally integrated the genes and transcripts identified by newly sequenced Iso-Seq data into the Gramene annotation results, as the basics to build homology relationships between 18 annotations. (Supplemental Table 1) In addition, a series of Iso-Seq and RNA-Seq data of multiple tissues from selected accessions (Supplemental Table 2) were collected and fully presented as baseline information in RGI, which included gene expression, full-length transcripts, and alternative splicing (AS) events. Details on data processing are described in supplemental methods.

**Figure 1.**
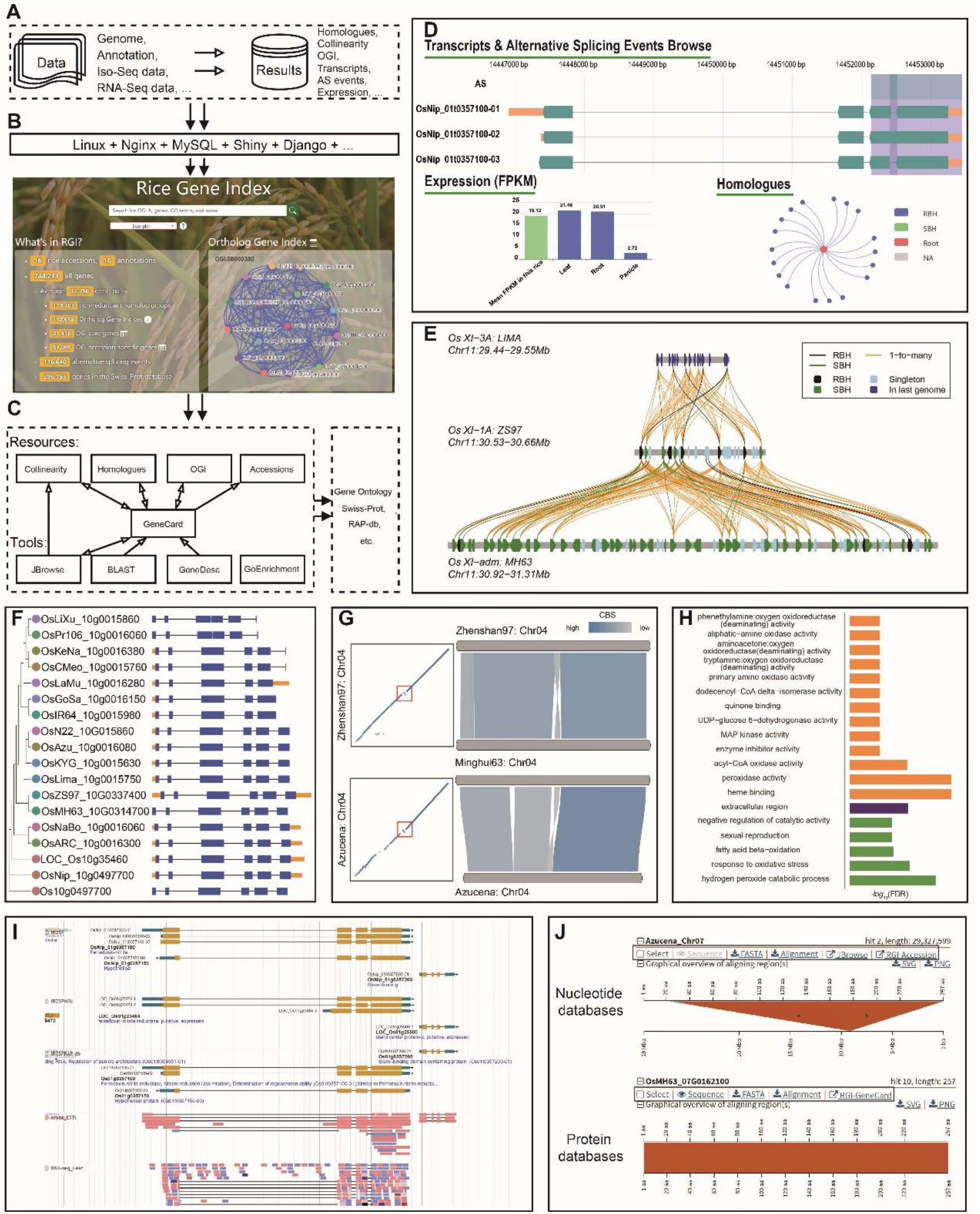
Design and online output example of RGI. (A-C) Overview of the construction of the Rice Gene Index (RGI). (A) The data layer demonstrates the source data and results provided in RGI. (B) The middleware layer shows the tools used to build the RGI website. (C) The architecture of RGI shows functions and tools. ‘GeneCard’ is a central module in RGI, which links to other modules via a network. (D) ‘GeneCard’ page shows *OsNiR*’s transcripts structure, AS events, Gene expression, and homologs. (E) ‘MicroColinearity’ module shows homologous relationships of a disease-resistant gene cluster on the local scale between the MH63 and ZS97 genomes. The black, green, and yellow lines represent RBH, SBH, and 1-to-many relationships, respectively. For genes, black, green, blue, and yellow represent genes with RBH, SBH, singleton, and 1-to-may relationships, respectively. (F) Gene tree in the ‘Homologues’ module. (G) ‘MacroCollinearity’ module shows the collinear blocks in chromosome 4 between ZS97 and MH63, and chromosome 4 between MH63 and Azucena. Colors indicate the collinear block scores. Inversions were highlighted by red triangle. (H) GO enrichment analysis. (I) *OsNiR’*s location in JBrowse. (J) The partial result of searching *GHD7*’s sequence in sequence databases of the whole genome (from 16 assemblies) and protein (from 18 annotations) by the ‘BLAST’ tool. The black triangles show the functions and links to other pages.

## Result

As the primary datasets in RGI, the genome annotations of 16 rice accessions contained an average of 41,346 genes, of which an average of 1,178 genes are supplemented by Iso-Seq data (Supplementary Table 3). The GeneTribe pipeline (Chen et al., 2020) identified an average of 33,350 gene pairs between annotations (Supplemental Figure 2), which classified ‘reciprocal best hits’ (RBHs), ‘single-side best hits’ (SBHs), ‘1-to-many’ hits or ‘singleton’ hits. By counting unique homolog gene groups, a total of 119,783 non-redundant gene groups were determined to represent the whole Asian rice gene set. To further unify the gene groups in *Oryza sativa*, we defined a unified and sustainable number — Ortholog Gene Index (OGI), which is a homolog group clustered by connected graph methods based on RBH relationships, with an updatable score that indicates its representativeness in all accessions. Of the 112,658 Ortholog Gene Indices, we classified them into 21,418 OGI core genes (19.01% of OGI) appearing in all rice accessions, 40,141 OGI dispensable genes, and 51,099 OGI accession-specific genes (Supplemental Figure 1A). And we found that the specific genes are younger and shorter (T-test, *p* = 2e-16) than core genes (Supplemental Information 1).

### Utility

The first objective of RGI is to logically organize and scientifically index all genes among rice accessions. RGI provides ‘GeneCard’ to show comprehensive information for individual genes with convenient links to other modules and outside databases on one page (Figure 1C). By entering a gene ID of rice, through the search box on the homepage, users may browse the ‘GeneCard’ page on 3 sections: 1) Basic information includes sequence, gene function, gene expression, links for accessing various modules and other databases, etc. (Supplemental Figure 4A). 2) ‘Transcripts’ exhibits graph and table of transcripts structure. In addition to the baseline expression analysis of all genes, 116,640 AS events at the transcriptome level were extensively revealed by the analysis of different groups (Supplemental Figure 4B, Supplemental Table 4). For example, two AS events were detected for *OsNiR* (OsNip_01g0357100), a critical gene that encodes nitrite reductase in nitrogen assimilation (Yu et al., 2021) (Figure 1D). Additionally, ‘Homologues’ lists all associated homologs of a gene across annotations through links graph and table ways. This section also shows the phylogenetic tree. Furthermore, RGI provides informative pages to show the association graph of genes in each Ortholog Gene Index (Supplemental Figure 4C).

Second, RGI provides three ways to search for relationships and comprehensive information for genes.

1. Through keyword-based searches, users can easily search OGI#, gene ID, gene symbol, Gene Ontology, or functional terms in the query box. If users search the famous gene *SD1* in RGI, 306 items will be returned with basic information, which could link to other modules or databases.
2. In the way of sequence-based searches, the classical ‘BLAST’ tool allows users to query amino acid or nucleotide sequences in sequence databases of the whole genome and protein. To easily access other modules, the tool returns gene ID linking to ‘GeneCard’ or chromosome location linking to ‘JBrowse’ when using the protein or nucleotide database, respectively.
3. For association-based searches, the ‘Homologues’ module allows users to query and connect the homologous genes through a given gene ID, which may obtain the homology relationship among annotations then build a homologous gene tree and display gene structures (Figure 1F), as well as multiple sequence alignments and positions on the chromosome of these homologs. For example, *OsTPP7* (LOC_Os09g20390), an anaerobic germination tolerance gene, was found to be absent in IR64 but present in other accessions by ‘Homologues’ (Supplemental Table 5) and manually verified the results. It indicates that IR64 has less tolerance to anaerobic germination (Yang et al., 2019).

Third, RGI can visualize the relationship of these annotated genes across accessions at local and global scales corresponding to two modules as follows:

1. At the local scale, the ‘MicroCollinearity’ module enables users to demonstrate genomic collinearities of a gene and its flanking genes in selected accessions (Figure 1E). The homologous relations among genomes help to investigate gene-based variations in the local regions of multiple accessions. Many genes encoding nucleotide-binding site leucine-rich repeat (NLR) proteins are found in the region close to the end of rice chromosome 11 long arm (Supplemental Figure 5) (Song *et al*., 2021), the collinearity comparison results detected by this module show that these NLR genes are significantly more abundant in MH63 than in other accessions, which potentially contribute to MH63’s superior resistance to rice diseases.
2. At the global scale, ‘MacroCollinearity’ helps users to explore collinearity between accessions and study rearrangements of rice genome at the whole-chromosome level. With this module, structure variations may be easily detected and the interactive tool ‘Dot Plot’ was embedded to show the collinearity details and links to associated genome loci on ‘JBrowse’. (Figure 1G). A useful module ‘GenePair’ is provided to visualize collinearity comparisons of ortholog gene pairs between two accessions on both global and local scales.

All information mentioned above is logically organized and seamlessly integrated by modules and tools in RGI. Four extra modules (‘JBrowse’ (Figure 1I), ‘GOEnrichment’ (Figure 1H), ‘GeneDescription’, and ‘Download’) were additionally integrated to enhance RGI’s serviceability (Supplemental Information 2). The technical details on RGI construction of RGI are described in Supplemental Information 3.

## Discussion

Although more than 100 chromosomal-level genomes of Asian rice have been published, most of the relevant databases focus on single genomes for specific domains (e.g., LncRNA, epigenomic, etc.) (Copetti et al., 2015; Xie et al., 2021; Zhang et al., 2021). Two ‘pan-genome’ databases have been published (i.e. RPAN (https://cgm.sjtu.edu.cn/3kricedb/index.php) provides data on individual rice accessions, and Rice RC (http://ricerc.sicau.edu.cn/RiceRC) has a focus on structure variants), while our RGI comprehensively creates and focuses on gene-level relationships across representative Asian rice accessions, establishes a standardized gene index for Asian rice, and provides richer search and visualization capabilities for the whole rice research community.

## DECLARATIONS

### Availability of data and materials

The datasets generated during and/or analyzed during the current study are available in https://riceome.hzau.edu.cn/. PacBio Iso-Seq raw data are available from NCBI under BioProject PRJNA760839. The RNA-Seq raw data are available from NCBI under BioProject PRJNA659864 and PRJNA5970706.

### Funding

This research was supported by Fundamental Research Funds for the Central Universities (2662020SKPY010) and the Major Project of Hubei Hongshan Laboratory (2022HSZD031) to J.Z.

### Authors’ contributions

R.A.W., W.G. and J.Z. designed and conceived the research. K.L.M. provided SSD seed, extracted DNA and RNA for genome and transcript sequencing (both by PacBio and Illumina) for 12 Asian rice. Z.Y., Y.Z., YU.Z. and M.L. performed the homologues and transcriptome analysis. Z.Y. and Y.C. built the database and managed the computing platforms. Z.Y., Y.C., Y.Z., Y.O., D.C., R.M., H.Z., W.X., K.L.M., R.A.W., W.G. and J.Z. wrote and edited the paper. All authors read and approved the final manuscript.

## Acknowledgments

We thank access to the annotation for the Magic 16 gene structure annotation from the Gramene project, specifically K. Chougule, Z. Lu, and D. Ware supported by USDA 8062-21000-041-00D. We sincerely thank the computing platform of the National Key Laboratory of Crop Genetic Improvement in HZAU for providing the computational resources. No conflict of interest declared.

## Supplemental Information

See the Supplementary Information.

### Figures

**Supplemental Figure 1.**
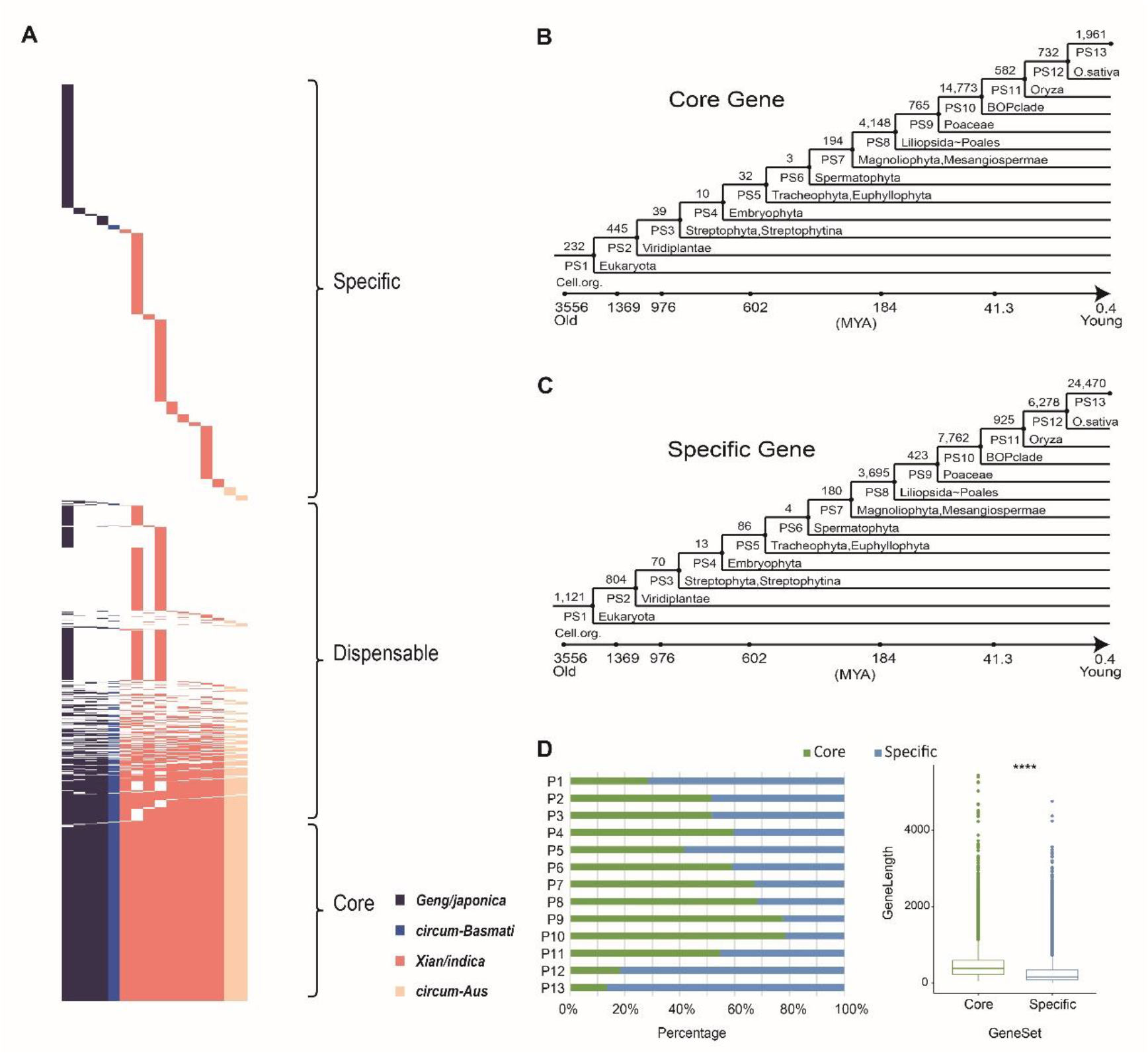
Gene presence/absence variations in 16 Asian rice accessions. (A) The core genes are present in all accessions, the dispensable genes are present in <16 of accessions, and the accession-specific genes are only present in one accession. The presence of genes is colorful (4 subpopulations in Asian rice: *Geng/japonica*, *Xian/indica*, *circum-Aus*, and *circum-Basmati*), and the absence of genes is white. (B and C) The numbers of core genes (B) and accession-specific genes (C) that emerged at different evolutionary times, from PS1 (single-cell organisms) to PS13 (*O. sativa*). (D) The age distribution (left) and gene length (right) of the core and genes. (See Methods)

**Supplemental Figure 2.**
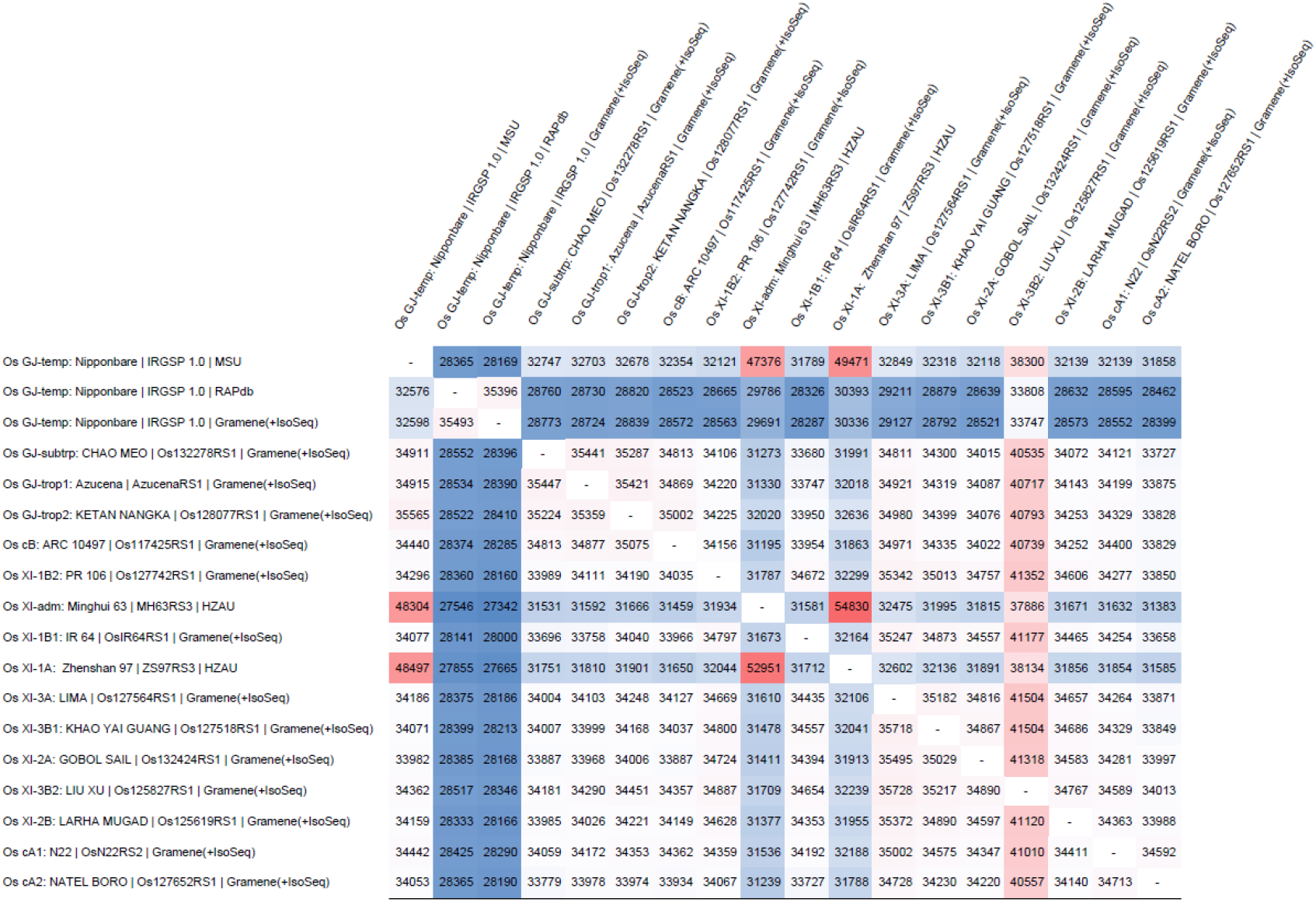
The 18 × 18 gene annotation matrix, which shows the number of 1-to-1 homologs, and demonstrates the homologous gene pairs. The color corresponds to the number of homologs. Red represents higher gene pairs number and blue represents lower gene pairs number.

**Supplemental Figure 3.**
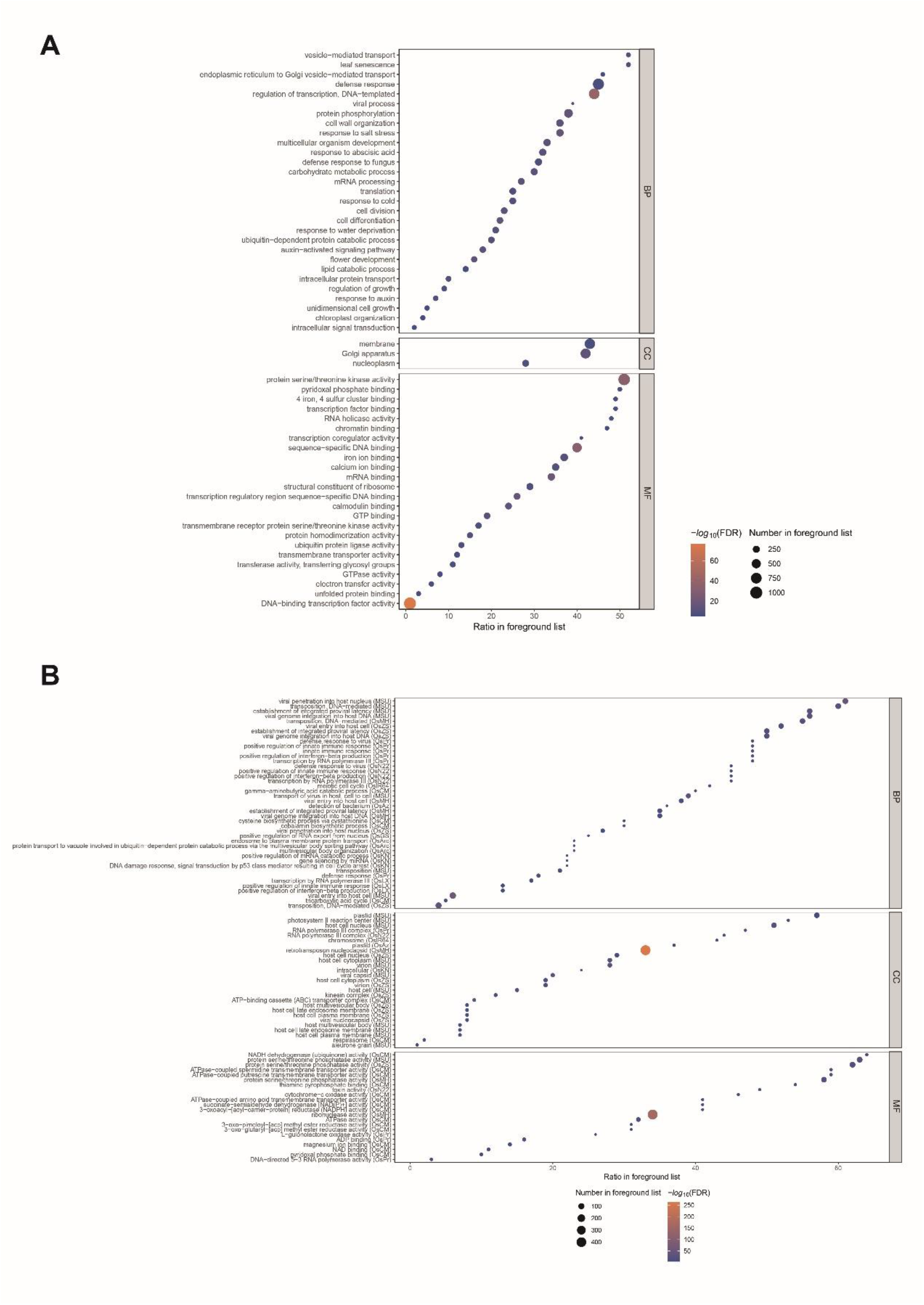
The bubble plot of the Gene Ontology enrichment analysis in (A) the core gene set and (B) the accession-specific gene set.

**Supplemental Figure 4.**
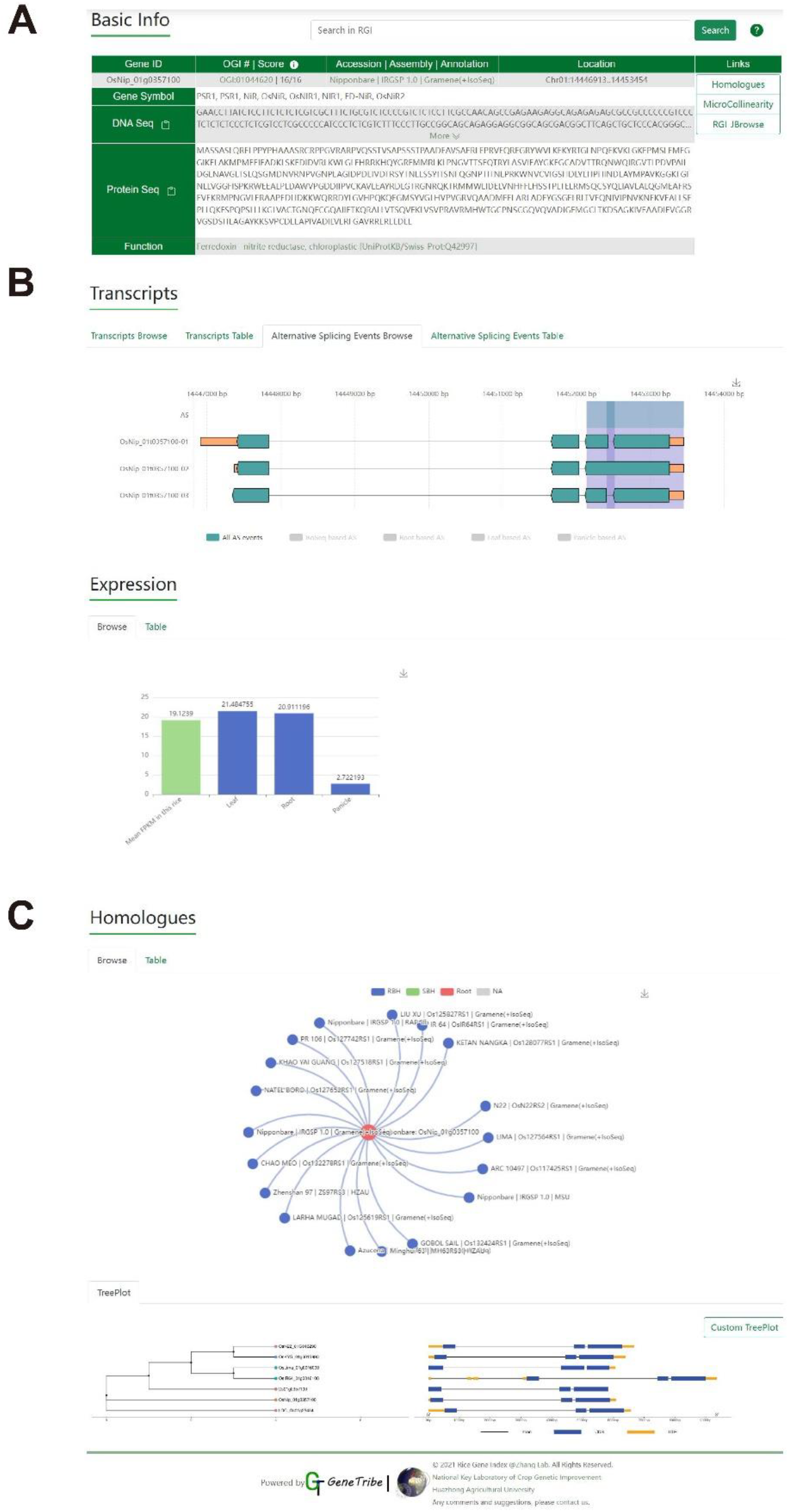
The ‘GeneCard’ page.

**Supplemental Figure 5.**
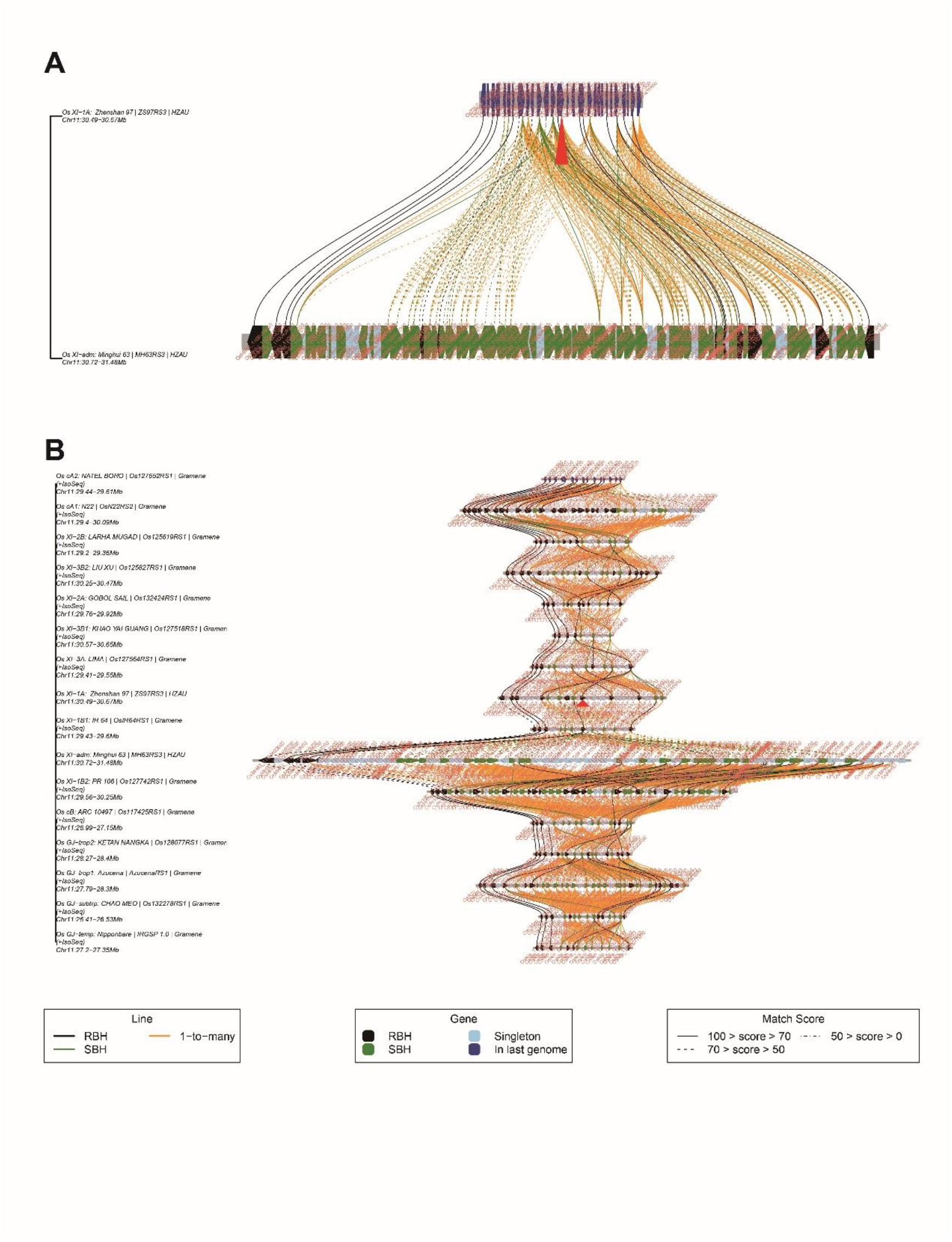
In previous studies, NLR genes were found to be highly duplicated in chromosome 11 of MH63 compared with other genomes. (A) displays the micro-collinearity of NLR genes in MH63 and ZS97 by searching NLR gene *OsZS97_11G0430400* in the ‘MicroCollinearity’ module. (B) displays the micro-collinearity of NLR genes in 16 varieties by searching NLR gene *OsZS97_11G0430400* in the ‘MicroCollinearity’ module. (A) and (B) show *OsZS97_11G0430400*’s 15 flanking genes.

### Supplementary Tables

**Supplemental Table 1.** Summary of RGI data composition.

| Supplemental Table 1. Summary of RGI data composition |  |  |  |  |
| --- | --- | --- | --- | --- |
| Accession Name | Subpopulations | Assembly | Annotation<sup>a</sup> | Locus Tag Prefix |
| Os GJ-temp: Nipponbare | Geng-japonica-temp | IRGSP1.0 | Gramene (+Isoseq) | OsNip_ |
|  |  |  | MSU | LOC_Os |
|  |  |  | RAP-db | Os |
| Os GJ-subtrp: CHAO MEO | Geng-japonica-trop1 | AzucenaRS1 | Gramene (+Isoseq) | OsAzu_ |
| Os GJ-trop1: Azucena | Geng-japonica-trop2 | Os128077RS1 | Gramene (+Isoseq) | OsKeNa_ |
| Os GJ-trop2: KETAN NANGKA | Geng-japonica-subtrp | Os132278RS1 | Gramene (+Isoseq) | OsCMeo_ |
| Os cB: ARC 10497 | circum-Basmati | Os117425RS1 | Gramene (+Isoseq) | OsARC_ |
| Os XI-1B2: PR 106 | Xian-indica-1B2 | Os127742RS1 | Gramene (+Isoseq) | OsPr106_ |
| Os XI-adm: MH63 | Xian-indica-adm | MH63RS3 | HZAU | OsMH63_ |
| Os XI-1B1: IR 64 | Xian-indica-1B1 | OsIR64RS1 | Gramene (+Isoseq) | OsIR64_ |
| Os XI-1A: ZS97 | Xian-indica-1A | ZS97RS3 | HZAU | OsZS97_ |
| Os XI-3A: LIMA | Xian-indica-3A | Os127564RS1 | Gramene (+Isoseq) | OsLima_ |
| Os XI-3B1: KHAO YAI GUANG | Xian-indica-3B1 | Os127518RS1 | Gramene (+Isoseq) | OsKYG_ |
| Os XI-2A: GOBOL SAIL | Xian-indica-2A | Os132424RS1 | Gramene (+Isoseq) | OsGoSa_ |
| Os XI-3B2: LIU XU | Xian-indica-3B2 | Os125827RS1 | Gramene (+Isoseq) | OsLiXu_ |
| Os XI-2B: LARHA MUGAD | Xian-indica-2B | Os125619RS1 | Gramene (+Isoseq) | OsLaMu_ |
| Os cA1: N22 | circum-Aus1 | OsN22RS2 | Gramene (+Isoseq) | OsN22_ |
| Os cA2: NATEL BORO | circum-Aus2 | Os127652RS1 | Gramene (+Isoseq) | OsNaBo_ |

a. this row is the annotation source, Gramene (+Isoseq) means *de novo* annotation attached transcripts from Iso-Seq data

**Supplemental Table 2.**
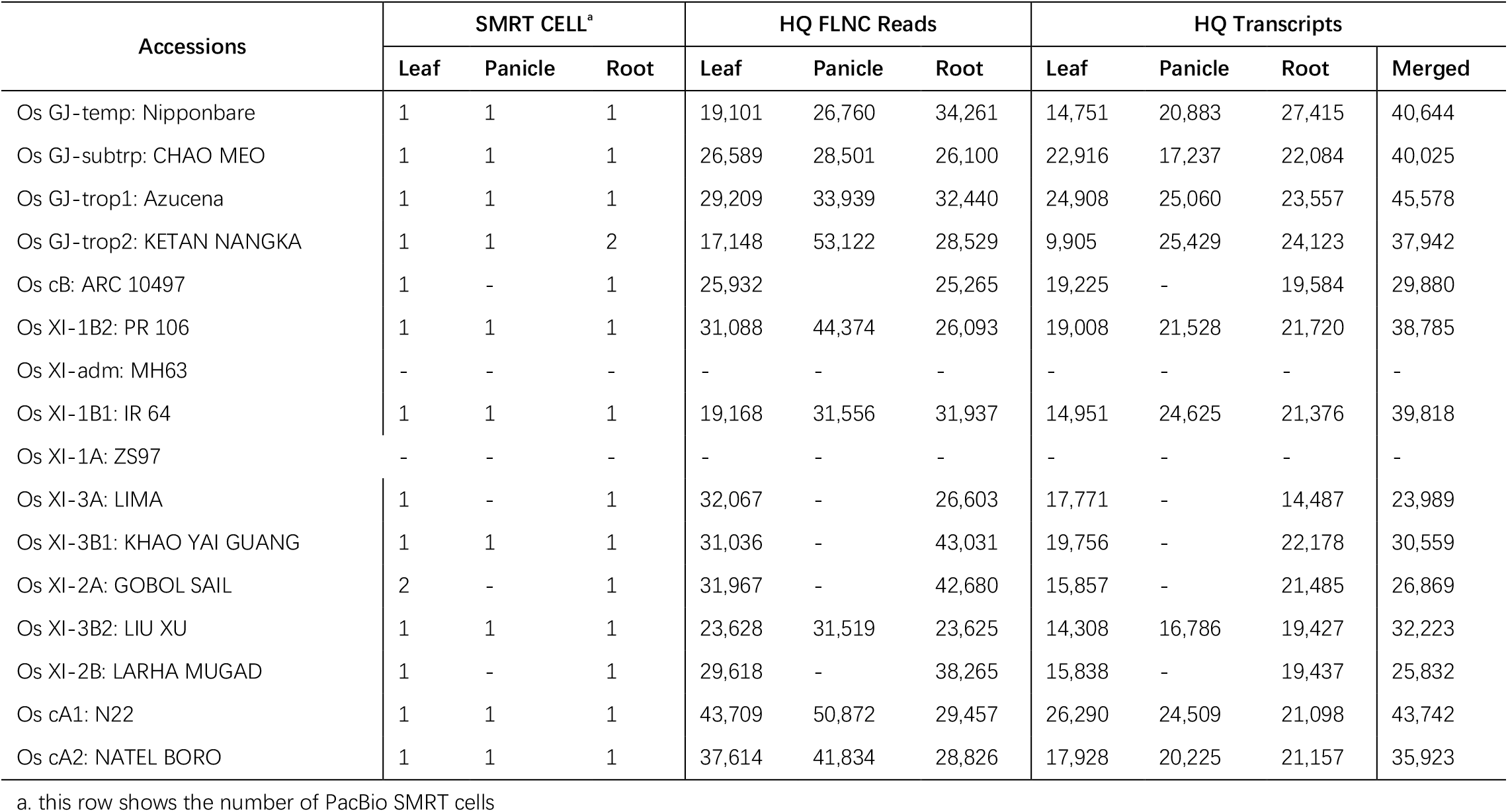
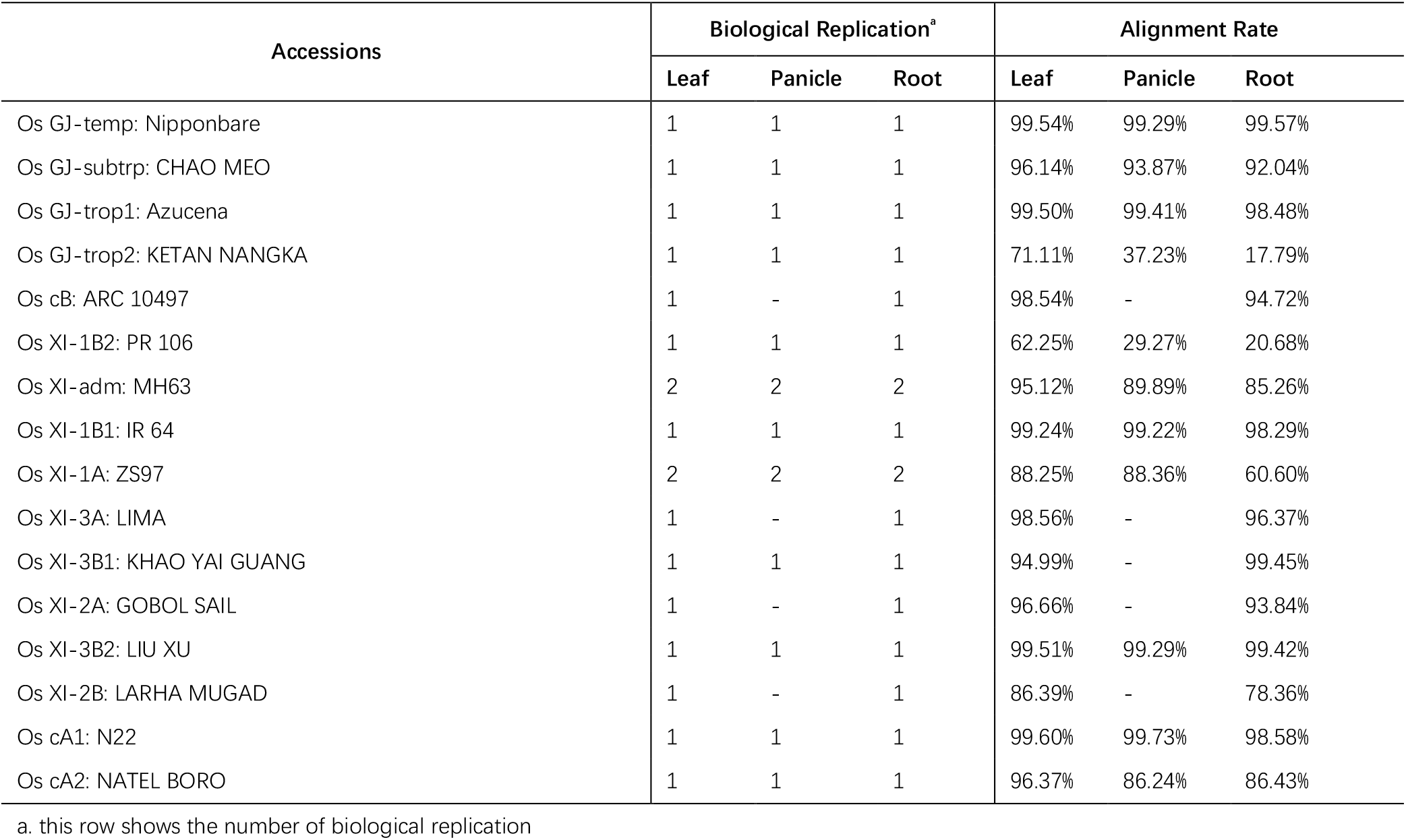
Iso-Seq and RNA-Seq data from multiple tissues (i.e., leaves, roots, and immature panicles) of 16 rice accessions.

**Supplemental Table 3.** Numbers of annotated genes in 16 rice accessions.

| Accession\|Assembly\|Annotation | Annotated Genes | Genes from Iso-Seq Data |
| --- | --- | --- |
| Os GJ-temp: Nipponbare \| IRGSP 1.0 \| MSU | 55,801 | - |
| Os GJ-temp: Nipponbare \| IRGSP 1.0 \| RAPdb | 37,859 | - |
| Os GJ-temp: Nipponbare \| IRGSP 1.0 \| Gramene(+IsoSeq) | 38,404 | 733 |
| Os GJ-subtrp: CHAO MEO \| Os132278RS1 \| Gramene(+IsoSeq) | 37,240 | 1,054 |
| Os GJ-trop1: Azucena \| AzucenaRS1 \| Gramene(+IsoSeq) | 36,882 | 1,079 |
| Os GJ-trop2: KETAN NANGKA \| Os128077RS1 \| Gramene(+IsoSeq) | 37,994 | 1,388 |
| Os cB: ARC 10497 \| Os117425RS1 \| Gramene(+IsoSeq) | 37,181 | 844 |
| Os XI-1B2: PR 106 \| Os127742RS1 \| Gramene(+IsoSeq) | 36,720 | 1,255 |
| Os XI-adm: Minghui 63 \| MH63RS3 \| HZAU | 59,903 | - |
| Os XI-1B1: IR 64 \| OsIR64RS1 \| Gramene(+IsoSeq) | 36,925 | 987 |
| Os XI-1A: Zhenshan 97 \| ZS97RS3 \| HZAU | 60,935 | - |
| Os XI-3A: LIMA \| Os127564RS1 \| Gramene(+IsoSeq) | 37,994 | 973 |
| Os XI-3B1: KHAO YAI GUANG \| Os127518RS1 \| Gramene(+IsoSeq) | 37,522 | 1,105 |
| Os XI-2A: GOBOL SAIL \| Os132424RS1 \| Gramene(+IsoSeq) | 36,467 | 1,810 |
| Os XI-3B2: LIU XU \| Os125827RS1 \| Gramene(+IsoSeq) | 44,942 | 1,183 |
| Os XI-2B: LARHA MUGAD \| Os125619RS1 \| Gramene(+IsoSeq) | 37,474 | 1,244 |
| Os cA1: N22 \| OsN22RS2 \| Gramene(+IsoSeq) | 37,598 | 1,280 |
| Os cA2: NATEL BORO \| Os127652RS1 \| Gramene(+IsoSeq) | 36,392 | 1,664 |

**Supplemental Table 4.** Summary of alternative splicing events in 16 rice accessions.

| Accession\|Assembly\|Annotation | A3 | A5 | AF | AL | MX | RI | SE | Total |
| --- | --- | --- | --- | --- | --- | --- | --- | --- |
| Os GJ-temp: Nipponbare \| IRGSP 1.0 \| Gramene(+IsoSeq) | 1,250 | 634 | 237 | 91 | 2 | 1,181 | 326 | 3,721 |
| Os GJ-subtrp: CHAO MEO \| Os132278RS1 \| Gramene(+IsoSeq) | 2,761 | 2,574 | 530 | 182 | 113 | 2,228 | 1,715 | 10,103 |
| Os GJ-trop1: Azucena \| AzucenaRS1 \| Gramene(+IsoSeq) | 2,735 | 2,583 | 575 | 187 | 100 | 2,257 | 1,719 | 10,156 |
| Os GJ-trop2: KETAN NANGKA \| Os128077RS1 \| Gramene(+IsoSeq) | 2,484 | 2,350 | 509 | 167 | 101 | 2,072 | 1,581 | 9,264 |
| Os cB: ARC 10497 \| Os117425RS1 \| Gramene(+IsoSeq) | 2,253 | 2,009 | 451 | 151 | 91 | 1,912 | 1,396 | 8,263 |
| Os XI-1B2: PR 106 \| Os127742RS1 \| Gramene(+IsoSeq) | 2,530 | 2,415 | 519 | 169 | 116 | 2,153 | 1,637 | 9,539 |
| Os XI-adm: Minghui 63 \| MH63RS3 \| HZAU | - | - | - | - | - | - | - | - |
| Os XI-1B1: IR 64 \| OsIR64RS1 \| Gramene(+IsoSeq) | 2,634 | 2,434 | 546 | 208 | 96 | 2,177 | 1,646 | 9,741 |
| Os XI-1A: Zhenshan 97 \| ZS97RS3 \| HZAU | - | - | - | - | - | - | - | - |
| Os XI-3A: LIMA \| Os127564RS1 \| Gramene(+IsoSeq) | 1,841 | 1,670 | 367 | 127 | 62 | 1,791 | 1,182 | 7,040 |
| Os XI-3B1: KHAO YAI GUANG \| Os127518RS1 \| Gramene(+IsoSeq) | 1,986 | 1,770 | 403 | 149 | 60 | 1,792 | 1,236 | 7,396 |
| Os XI-2A: GOBOL SAIL \| Os132424RS1 \| Gramene(+IsoSeq) | 1,563 | 1,355 | 351 | 115 | 45 | 1,450 | 940 | 5,819 |
| Os XI-3B2: LIU XU \| Os125827RS1 \| Gramene(+IsoSeq) | 2,537 | 2,424 | 493 | 168 | 97 | 1,981 | 1,683 | 9,383 |
| Os XI-2B: LARHA MUGAD \| Os125619RS1 \| Gramene(+IsoSeq) | 1,803 | 1,588 | 377 | 154 | 57 | 1,611 | 1,129 | 6,719 |
| Os cA1: N22 \| OsN22RS2 \| Gramene(+IsoSeq) | 2,742 | 2,548 | 514 | 937 | 108 | 2,464 | 1,705 | 11,018 |
| Os cA2: NATEL BORO \| Os127652RS1 \| Gramene(+IsoSeq) | 2,281 | 2,198 | 474 | 156 | 104 | 1,796 | 1,469 | 8,478 |

**Supplemental Table 5.** Homologues of gene *LOC_Os11g29290* in 16 accessions. The result was produced by the ‘Homologues’ module and checked manually.

| Accession\|Assembly\|Annotation | Homologous | Type |
| --- | --- | --- |
| Os GJ-temp: Nipponbare \| IRGSP 1.0 \| MSU | LOC_Os11g29290 | RBH |
| Os GJ-temp: Nipponbare \| IRGSP 1.0 \| RAPdb | Os11g0483000 | RBH |
| Os GJ-temp: Nipponbare \| IRGSP 1.0 \| Gramene(+IsoSeq) | OsNip_11g0483000 | RBH |
| Os GJ-subtrp: CHAO MEO \| Os132278RS1 \| Gramene(+IsoSeq) | OsCMeo_11g0013950 | RBH |
| Os GJ-trop1: Azucena \| AzucenaRS1 \| Gramene(+IsoSeq) | OsAzu_11g0014030 | RBH |
| Os GJ-trop2: KETAN NANGKA \| Os128077RS1 \| Gramene(+IsoSeq) | OsKeNa_11g0013990 | RBH |
| Os cB: ARC 10497 \| Os117425RS1 \| Gramene(+IsoSeq) | OsARC_11g0013770 | RBH |
| Os XI-1B2: PR 106 \| Os127742RS1 \| Gramene(+IsoSeq) | OsPr106_11g0014050 | RBH |
| Os XI-adm: Minghui 63 \| MH63RS3 \| HZAU | OsMH63_11G0261700 | RBH |
| Os XI-1B1: IR 64 \| OsIR64RS1 \| Gramene(+IsoSeq) | NA | NA |
| Os XI-1A: Zhenshan 97 \| ZS97RS3 \| HZAU | OsZS97_11G0273800 | RBH |
| Os XI-3A: LIMA \| Os127564RS1 \| Gramene(+IsoSeq) | OsLima_11g0014120 | RBH |
| Os XI-3B1: KHAO YAI GUANG \| Os127518RS1 \| Gramene(+IsoSeq) | OsKYG_11g0014170 | RBH |
| Os XI-2A: GOBOL SAIL \| Os132424RS1 \| Gramene(+IsoSeq) | OsGoSa_11g0013820 | RBH |
| Os XI-3B2: LIU XU \| Os125827RS1 \| Gramene(+IsoSeq) | OsLiXu_11g0013660 | RBH |
| Os XI-2B: LARHA MUGAD \| Os125619RS1 \| Gramene(+IsoSeq) | OsLaMu_11g0013960 | RBH |
| Os cA1: N22 \| OsN22RS2 \| Gramene(+IsoSeq) | OsN22_11G013780 | RBH |
| Os cA2: NATEL BORO \| Os127652RS1 \| Gramene(+IsoSeq) | OsNaBo_11g0014000 | RBH |

## Notes

https://riceome.hzau.edu.cn/

### Competing Interest Statement

The authors have declared no competing interest.

