## SupplementaryInformation for "Rice Gene Index (RGI): a comprehensive pan-genome database for comparative and functional genomics of Asian rice"

### Supplemental Information

#### Supplemental Information 1 – Functional annotation of OGI genes across species

To gain insights into the evolutionary history of OGI genes, we aligned their protein sequences to the NR database partitioned into 13 taxonomic levels (Wang et al., 2018). We found that new genes explosively emerged with the appearance of *Poaceae* (PS10) and *O. sativa* (PS13) respectively (Supplemental Figures 1B, C) and the specific genes are younger and shorter (T-test, $p$ = 2e-16) than core genes (Supplemental Figure 1D). By functional analysis with InterPro domains screened for the gene set, we found that ~88.35% of coding genes contained InterPro domains in the core gene set, a much higher proportion than that in the specific gene set (~35.63%), implying that accession-specific genes may be new genes or pseudogenes. Furthermore, core genes are enriched in essential functions for development, regulation of transcription, DNA-binding transcription factor activity, and so on (using Gene Ontology, GO) (Supplemental Figure 3A), whereas accession-specific genes are enriched in biotic responses like viral penetration into host nucleus (Supplemental Figure 3B).

#### Supplemental Information 2 – Extra useful functional applications

The 'JBrowse' is deployed to present various data tracks like coding and non-coding annotations, transcriptome data, etc. (Figure 1I). 'GOEnrichment' (Figure 1H) provides GO enrichment analysis, 'GeneDescription' may retrieve gene functions by batches, 'OGI' displays the topological structure of each Ortholog Gene Index, and 'Download' supplies the genomes and annotations of 16 rice accessions.

#### Supplemental Information 3 – System construction

RGI is hosted on a Linux operation system and Nginx web server (https://www.nginx.com) (Figure 1B). All featured data (e.g., homologs, collinearity blocks, transcripts, AS events, and gene functions) of RGI were organized and stored in the MySQL database (http://www.mysql.com). The website was constructed by using Shiny (https://shiny.rstudio.com) and Django (https://www.djangoproject.com) for the web framework, Bootstrap (https://getbootstrap.com) for the front-end design, and Echarts (https://echarts.apache.org) for data visualization.

### Methods

**Iso-Seq data processing** Iso-Seq data were sequenced from leaves and roots in 14 accessions, and panicles in 9 accessions. The leaves and roots data were processed to high quality (>98%) reads from PacBio subreads raw data with -ccs_max_length 15000 -ccs_min_length 50 -ccs_polish false -hq_cutoff 0.98 -Maximum_Fuzzy_Junction_Difference 5 -Minimum_Mapped_Concordance 95 -Minimum_Mapped_Coverage 99 -Minimum_Mapped_Length 50 -Require_and_Trim_Poly(A)_Tail true -Run_Clustering true, using SMRTLink 9.0.0.92188 from Pacific Biosciences. The panicles data were sequenced in a mixed pool (9 samples), and the data were processed to circular consensus sequence (CCS) from PacBio subreads raw data with -Maximum_CCS_Read_Length 50000 -Minimum_CCS_Read_Length 10 -Minimum_Number_of_Passes 3 -Minimum_Predicted_Accuracy 0.98, using SMRTLink 8.0.0.80529. Then, we split 9 sample data by barcode and generated high quality (>98%) reads, using the standard IsoSeqv3 (https://github.com/PacificBiosciences/IsoSeq) pipeline. All high-quality (>98%) reads were mapped to the respective genome using minimap2 (v2.17) (Li, 2018). TAMA (Kuo et al., 2020) with -x no_cap -i 0.95 was used to collapse redundant transcripts. Furthermore, TAMA was used to merge the transcripts between tissues and those from CDS prediction.

**RNA-Seq data processing**

RNA-Seq data were sequenced from multiple tissues (i.e., leaves, roots, and immature panicles) in 16 accessions. The RNA-Seq data was mapped to respective genomes using HISAT2 (v2.1.0) (Kim et al., 2019) with default parameters. The gene expression was calculated from the alignment results using StringTie (v1.3.4d) (Pertea et al., 2015).

**Annotation reconstruction**

Gene annotations were reconstructed by *de novo* annotation and transcripts from Iso-Seq data. Based on the *de novo* annotation, we identify the new genes and transcripts from Iso-Seq data by comparing them in each variety using SQANTI3 (Tardaguila et al., 2018). Based on the *de novo* annotation gene ID, we inserted the new genes and transcripts to *de novo* annotations by gene locations using python scripts.

#### Homologs identification

Homologs were identified using reconstructed annotations. Proteins translated by the longest transcript in gene were used to identify homologs. Then, GeneTribe (Chen et al., 2020) was used to perform the homology inference, which combined sequence similarity and collinearity block information, and generated homology relationships including RBH, SBH, 1-to-many, and singletons.

#### Ortholog Gene Index establishment

First, we removed redundant of the homologous gene groups across 18 annotations to obtain 119,783 non-redundant homologous gene groups. Furthermore, using the above data, homologous gene groups were clustered with the connected graph algorithm to obtain 112,658 Ortholog Gene Indices (OGI). Furthermore, using the above data, homologous gene groups were clustered with the connected graph algorithm (McColl et al., 1986) to obtain 112,658 Ortholog Gene Indices (OGI). If two gene groups have overlapped genes, they will be clustered to a new gene group by removing the redundant homologous genes. And we iterated the step until there were no connected homologous relationships found between any groups. The indices were then named with the following rules: the first two digits are the chromosome number containing the most genes in each OGI, and the last six digits represent the order of genes on the genome (Using the order of genes in MH63 as a reference to determine the basic order of OGI, and for OGI not containing MH63 gene, query the position of MH63 gene in the OGI where its upstream gene is located, make insertion, and finally rearrange the order at intervals of 10.). For each OGI, a score was calculated and assigned with the number of involved accessions in each OGI divided by the total number of all accessions, and will be updated as the data increases.

**Gene ages identification**

Gene ages were identified by previous methods (Wang *et al.*, 2018). We downloaded the NR database (https://ftp.ncbi.nlm.nih.gov/blast/db/FASTA/) from NCBI (13 October 2020) and classified the protein sequence to 13 taxonomic levels (PS1 [Cellular organisms]; PS2 [Eukaryota]; PS3 [Viridiplantae]; PS4 [Streptophyta, Streptophytina]; PS5 [Embryophyta]; PS6 [Tracheophyta, Euphyllophyta]; PS7 [Spermatophyta]; PS8 [Magnoliophyta, Mesangiospermae]; PS9 [Liliopsida, Petrosaviidae, Commelinids, Poales]; PS10 [Poaceae]; PS11 [BOP clade]; PS12 [Oryzoideae, Oryzeae, Oryza]; and PS13 [Oryza.sativa]) based on NCBI taxonomy. OGI core gene set and specific gene set were translated into proteins and were aligned to the 13 databases using BLASTP (2.11.0+) (Camacho et al., 2009) with E-value < 1e-5 and identity > 30%. The age of a gene was considered as the taxonomic level of the oldest aligned protein.

**Gene function annotation**

Proteins from annotation were mapped to the Swiss-Prot database with -evalue 1e-10 using BLAST (2.11.0+) (Camacho *et al.*, 2009). The relationships between genes and Gene Ontology (GO) terms were extracted from the BLAST result by python scripts. We annotated the pan gene sets by InterProScan (v5.55-88.0) (Jones et al., 2014) with -dp -f tsv, and screened by evalue < 1E-10.

**Alternative splicing events identification**

SUPPA2 (2.3) (Trincado et al., 2018) was used to identify AS events for all transcripts, transcripts from Iso-Seq data and transcripts from various tissues (i.e., leaves, roots, and panicles). The AS events were classified into various types such as intron retention, exon skipping, alternative donor site, alternative accepter site, and alternative position.

**Phylogenetic analysis**

We first performed the multiple sequence alignment in homolog groups (longest protein of each gene) by mafft (7.475) (Katoh and Standley, 2013) with –auto –treeout –maxiterate 1000 –thread 8 –quiet –inputorder. Second, the result were trimmed by trimAl (1.4.1) (Capella-Gutiérrez et al., 2009) with -automated1. Finally, we used IQ-TREE (1.6.12) (Nguyen et al., 2014) to generate the phylogenetic tree with default parameters and show the tree by ggtree (Yu et al., 2017).

#### Other integrated Tools

The 'BLAST' tool was based on Sequenceserver (Priyam et al., 2019), and the 'JBrowse' was adapted from Buels, et al (2016).
